# Buffering Updates Enables Efficient Dynamic de Bruijn Graphs

**DOI:** 10.1101/2021.03.16.435535

**Authors:** Jarno Alanko, Bahar Alipanahi, Jonathen Settle, Christina Boucher, Travis Gagie

## Abstract

**Motivation:** The de Bruijn graph has become a ubiquitous graph model for biological data ever since its initial introduction in the late 1990s. It has been used for a variety of purposes including genome assembly (Zerbino and Birney, 2008; Bankevich et al., 2012; Peng et al., 2012), variant detection (Alipanahi et al., 2020b; Iqbal et al., 2012), and storage of assembled genomes (Chikhi et al., 2016). For this reason, there have been over a dozen methods for building and representing the de Bruijn graph and its variants in a space and time efficient manner.

**Results:** With the exception of a few data structures (Muggli et al., 2019; Holley and Melsted, 2020; Crawford et al., 2018), compressed and compact de Bruijn graphs do not allow for the graph to be efficiently updated, meaning that data can be be added or deleted. The most recent compressed dynamic de Bruijn graph (Alipanahi et al., 2020a), relies on dynamic bit vectors which are slow in theory and practice. To address this shortcoming, we present a compressed dynamic de Bruijn graph that removes the necessity of dynamic bit vectors by buffering data that should be added or removed from the graph. We implement our method, which we refer to as BufBOSS, and compare its performance to Bifrost, DynamicBOSS, and FDBG. Our experiments demonstrate that BufBOSS achieves attractive trade-offs compared to other tools in terms of time, memory and disk, and has the best deletion performance by an order of magnitude.

## 1. Introduction

Analyzing population level data has posed significant algorithmic challenges as the amount of the data has risen steadily in the past decade or more. For example, the 1,000 Genomes Project was concluded in 2012 (McVean et al.), and more recently, the 100K Project concluded in 2018 (Turnbull et al.). The Meta-Sub project (Danko et al., 2021), which collects metagenomic samples from subway systems across the world began in 2016, is now in its fifth year. One of the main goals of these projects is to identify rare variants – including but not limited to single nucleotide polymorphisms (SNPs), insertions, and deletions – within the population, and to attribute them to physiological or disease outcomes. One method of identifying these variants that receives a significant amount of attention is the construction and analysis of the *colored de Bruijn graph*.

To define the colored de Bruijn graph correctly it is useful to first define the de Bruijn graph in a constructive manner as follows. Given a set of sequence reads *R* and integer *k*, the first step is to identify all unique *k*-length substrings (*k*-mers) occurring in *R*, then create a directed edge for each unique *k*-mer with node labels being the (*k* − 1)-length prefix and the (*k* − 1)-length suffix of the *k*-mer, and lastly, after all directed edges have been created, glue all nodes that have the same label. The original purpose of the de Bruijn graph was to assemble single genomes (Pevzner et al., 2001), however, the definition and purpose has expanded to the linked de Bruijn graph (Turner et al., 2018), positional de Bruijn graph (Cameron et al., 2017; Ronen et al., 2012; Alipanahi et al., 2017) and paired de Bruijn graph (Medvedev et al., 2011) to name a few. The colored variant of the de Bruijn graph is arguably the most well-studied. It is constructed using the sequence data of a population of genomes, rather than a single genome. Here, we assume we have *d* sets of sequence reads that are denoted as *R*_1_, *R*_2_, .., *R*_*d*_, and we assign a unique color for each these *d* sets. Using this assignment of colors, each edge (and node) is colored with color *c*_*i*_ if and only if the associated *k*-mer ((*k* − 1)-mer) is contained in *R*_*i*_. In this way, each edge (or node) is assigned one or more colors which signify the sets of sequence reads which contain the corresponding *k*-mer ((*k* − 1)-mer). The colored de Bruijn graph can then be seen as a de Bruijn graph constructed from the union of the *k*-mers from *R*_1_, .., *R*_*d*_, and a binary matrix *C* that stores the colors of each *k*-mers, i.e., *C*[*i, j*] = 1 if the *i*-th *k*-mer is contained in *R*_*j*_. Thus, by traversing the graph and finding shared and divergent paths, rare genetic variants occurring in the population can be identified.

Iqbal et al. (2012) introduced the colored de Bruijn graph and gave the first implementation, which was in turn used to analyze the 1,000 Genomes Project data. This initial implementation was not space efficient but it motivated the need for implementations that could handle population-level datasets. Hence, in the past couple years there has been an explosion in the interest in space-efficient colored de Bruijn graphs. Vari (Muggli et al., 2017) and Rainbowfish (Almodaresi et al., 2017), were the first space-efficient colored de Bruijn graphs methods. Both recognize that the colored de Bruijn graph can be made space-efficient by storing a succinct de Bruijn graph built from the union of all sets of sequence reads (i.e., *R*_1_ ∪*R*_2_ ∪… ∪*R*_*d*_), and a compressed color matrix. Bloom Filter Trie (BFT) (Holley et al., 2015) was another early method for compactly storing the colored de Bruijn graph that uses Bloom filters to store the de Bruijn graph. These initial developments were followed by many other methods, including Mantis (Pandey et al., 2018), and Bifrost (Holley and Melsted, 2020).

Yet, several public consortium projects are not only significantly large but are also continually evolving, including GenomeTrakr (Allard et al., 2016) and MetaSub (Danko et al., 2021). For this reason, there is growing interest in dynamic, space-efficient colored de Bruijn graphs that can also evolve with the data – meaning that they allow addition and deletion of data. Given that there are methods for efficiently storing the color matrix in manner that is dynamic, one of the remaining challenges is how to store the de Bruijn graph in a manner that is mutable but also space and time efficient. This is challenging since mutability and compressibility are contradictory in nature. Even allowing for the addition of data into a compressed version of the de Bruijn graph requires some algorithmic cleverness. For example, VariMerge (Muggli et al., 2019) and Bifrost (Holley and Melsted, 2020) allow addition of new data but cannot perform deletion. VariMerge enables addition into a colored de Bruijn graph that is stored using Vari by implementing an algorithm that merges it with another colored de Bruijn graph that is also stored using Vari. The merge is done without any decompression for space-efficiency. Bifrost first constructs the de Bruijn graph using Bloom filters, then extracts all non-overlapping and non-branching paths in the graph (unitigs) and indexes all the unitigs using minimizers. It performs addition by rebuilding the de Bruijn graph from the minimizer representation, building a new graph for the additions, and then – similar to VariMerge – merging these two graphs succinctly. We refer the reader to Marchet et al. for a more thorough explanation of some of these key concepts, including minimizers and Bloom filters.

Similarly, there are only two compressed data structures for storing the de Bruijn graph that allows for both addition and deletion of data; these are FDBG (Crawford et al., 2018) and DynamicBOSS (Alipanahi et al., 2020a). FDBG is an implementation of the hash-based data structure of Belazzougui et al. (2016a). Dynamic-BOSS is more closely related to the work in this paper. It adapts the de Bruijn graph representation of Bowe et al. (2012), which represents a de Bruijn graph using Burrows Wheeler Transform (BWT). Although DynamicBOSS is capable of handling large datasets and is fully dynamic, it is slow due to the reliance on dynamic bit vector libraries, i.e., the library of Prezza (2017). Although dynamic bit vectors are being improved, we think it would be better to avoid their use and implement dynamism using static data structures instead. In Table 1, we give an overview of the basic de Bruijn graph data structures and the operations that they support.

**Table 1:** An overview of recent de Bruijn graph implementations and their attributes. In theory, BFT is capable of addition of data but it is no longer supported or functional (Holley, 2019).

|  | BufBOSS | DynamicBOSS | FDBG | Bifrost | Pufferfish | BFT | Mantis |
| --- | --- | --- | --- | --- | --- | --- | --- |
| Addition | Yes | Yes | Yes | Yes | No | No | No |
| Deletion | Yes | Yes | Yes | No | No | No | No |

Our contribution is a dynamic, space-efficient de Bruijn graph representation that eliminates the need for dynamic bit vectors. We implement dynamism using the combination of a static, space-efficient representation of the de Bruijn graph, and two auxiliary data structures, which buffer data that is to be added and deleted. As requests for additions or deletions are made, the corresponding buffers are updated, and after a prescribed number of additions or deletions, the static de Bruijn graph is rebuilt. This amortizes the cost of dynamic operations over the cost of rebuilding from time to time. Traversal or use of the graph takes into account the static de Bruijn graph as well as the addition and deletion buffer, which thus makes the static nature of the underlying data structure opaque. We describe how to: (a) buffer the data that is to be added or deleted, (b) support graph traversal taking into account the updates in the buffer (c) merge the updates to the graph efficiently.

We compare our method to existing dynamic de Bruijn graphs – namely Bifrost (Holley and Melsted, 2020), FDBG (Crawford et al., 2018), and Dynamic-BOSS (Alipanahi et al., 2020a). The evaluation criteria are the memory and time needed to construct the data structure, the size of the data structure on disk, the time required to add and delete data from the graph, and the time required to perform look-up queries of *k*-mers. In addition to these existing methods, we also implement and compare against a modified version of FDBG that we refer to as FDBG-RecSplit, where RecSplit (Esposito et al., 2020b) is used for minimal perfect hashing. RecSplit is the most recent minimal perfect hash implementation and thus, shows to have superior computational performance against competing methods.

BufBOSS was clearly superior to all methods that support addition and deletion (i.e., DynamicBOSS and FDBG) in both memory and time on all large datasets. Bifrost was the closest competitor but does not support deletion and our method was more performant than Bifrost in construction on large metagenomic datasets. In particular, our results show that BufBOSS was up to five times faster and used up to 30% less memory to construct than the closest competitor (Bifrost). The time required to add new sequences to BufBOSS was the second fastest in the competition, losing only to Bifrost by a factor of two, but beating the other tools by more than a factor of ten. The peak space during additions was two times larger than Bifrost, but we can vary a time-space trade-off parameter to push our space to slightly below Bifrost with a cost of slowing down the additions by a factor of five. With this setting, we are still faster than the rest of the tools, with a space that is smaller by a factor of eight or more. Lastly, we note the performance of deletion was the best out of all methods that support deletion, i.e., by a factor of 26 in time, and 13 in memory.

## 2. Preliminaries

In this section, we briefly go over some of the basic terminology and definitions that will be used throughout this paper.

### 2.1. Basic definitions

A string *X*, is represented as a sequence of characters: *X*[1..*n*] = *X*[1]*X*[2] … *X*[*n*], where *n* is the length of the string. Each character will be drawn from an ordered alphabet Σ of size σ. *X*[*i*.. *j*] refers to the substring of *X* given by *X*[*i*]*X*[*i* + 1] … *X*[*j*], where 1 ≤ *i* ≤ *j* ≤ *n*. Using this notation, a suffix is a substring with *j* = *n*, while a prefix is a substring with *i* = 1. An empty string is denoted with ε, and it is considered to be a prefix and a suffix of any string. A string of length *k* is called a *k*-mer. The lexicographic order of two strings *X* and *Y* is defined by the order of the characters at the first mismatching position from the left. If no mismatch exists – i.e., one string is a prefix of the other – the prefix is defined to be smaller. The *colexicographic* order of strings is the same but characters are compared from right to left, and in case of no mismatch, *X* is smaller than *Y* iff *X* is a suffix of *Y*.

In this paper, we use the *edge-centric* definition of de Bruijn graphs. In this definition, the graph is such that the nodes are all the distinct (*k* − 1)-mers of the input strings. There is an edge from *u* ((*k* − 1)-mer) to *v* ((*k* − 1)-mer) iff there exists a *k*-mer in some input string that is prefixed by *v* and suffixed by *u*.

To mitigate confusion on whether *k* refers to the length of edges or nodes, we call the (*k* − 1)-mers represented by the nodes *nodemers* and the *k*-mers represented by the edges *edgemers*. We denote the (*k* − 1)-mer represented by a node *v* with 𝓁 (*v*) and the *k*-mer represented by an edge *e* with 𝓁 (*e*). A nodemer or an edgemer from the DNA alphabet ACGT is said to be *canonical* if it is equal to its reverse complement.

### 2.2. Overview of BOSS

The BOSS data structure (Bowe et al., 2012) is a generalization of the classic FM-index (Ferragina and Manzini, 2005) to de Bruijn graphs. It is a special case of the more general *Wheeler graph index* (Gagie et al., 2017). Here, we will describe the BOSS data structure in terms of the more general Wheeler graph index, as our implementation is based on that representation.

The Wheeler graph index for a de Bruijn graph *G* is based on a larger graph *W* that contains *G* as a subgraph. The edges of *W* are labeled by single characters, and the nodes are unlabeled. Edges are labeled by the last characters of the corresponding edgemers in *G*. We denote the single-character label of an edge *e* in *W* with *L*(*e*) to distinguish from the notation 𝓁 (*e*) that refers to the *k*-mer corresponding to *e*.

This construction guarantees that all incoming paths of length (*k* − 1) to a node *v* spell the same string. Thus, one can walk back (*k* − 1) nodes in the graph to extract the (*k* − 1)-mer represented by a node. However, in subgraph *G* alone, some nodes might not have an incoming path of length (*k* − 1). To amend this, we add for every node *v* with indegree 0 a new path of *k* nodes, each representing a proper prefix of the (*k* − 1)-mer represented by *v*, all the way to the empty prefix ε. The added edges are labeled by the last character of the corresponding prefix of the destination node. If the same prefix is added for two different nodes, the nodes representing those prefixes are glued together. These added nodes and edges are called *dummy nodes* and *dummy edges*. The dummy nodes and edges form a tree starting from the node of the empty string, such that the leaves of the tree have outgoing edges to nodes of subgraph *G*. The notations 𝓁 (*v*) and 𝓁 (*e*) are extended for dummy nodes *v* and dummy edges *e* to denote the prefix represented by the node or the edge. The notation *L*(*e*) again denotes the last character of 𝓁 (*e*).

The Wheeler index on *W* is based on the colexicographical order ≺ on the nodes and edges of *W*. For nodes *u* and *v*, it is defined that *u* ≺ *v* iff 𝓁 (*u*) is colexicographically smaller than 𝓁 (*v*). The edges *e* are ordered colexicographically among themselves in the same way by 𝓁 (*e*). The Wheeler index consists of three main components: (1) a string EBWT, which is the concatenation of all edge labels of *W* sorted by the order of the node at the origin of the edge, with ties broken arbitrarily (2) a bit vector encoding the outdegrees of all nodes of *W* in the colexicographic order (3) a bit vector encoding the indegrees of all nodes of *W* in the colexicographic order. Remarkably, these data structures define the graph completely (Gagie et al., 2017).

Figure 1 shows an example of a Wheeler graph. The EBWT is illustrated with vertical separators showing where the node at the origin changes. These separators are for illustration purposes only and are not included in the EBWT string. The sequence of outdegrees in colexicographic order is 1,2,1,0,0,1,1,1,1,1,2,1,1, whereas the sequence of indegrees is 0,1,1,1,2,1,1,1,1,1,1,1,1. The Wheeler graph index represents these by encoding degree *d* with bit string 1· 0^*d*^, and concatenating the representations. The concatenated representation of outdegrees is denoted with *O* and the concatenated representation of indegrees with *I*.

**Figure 1:**
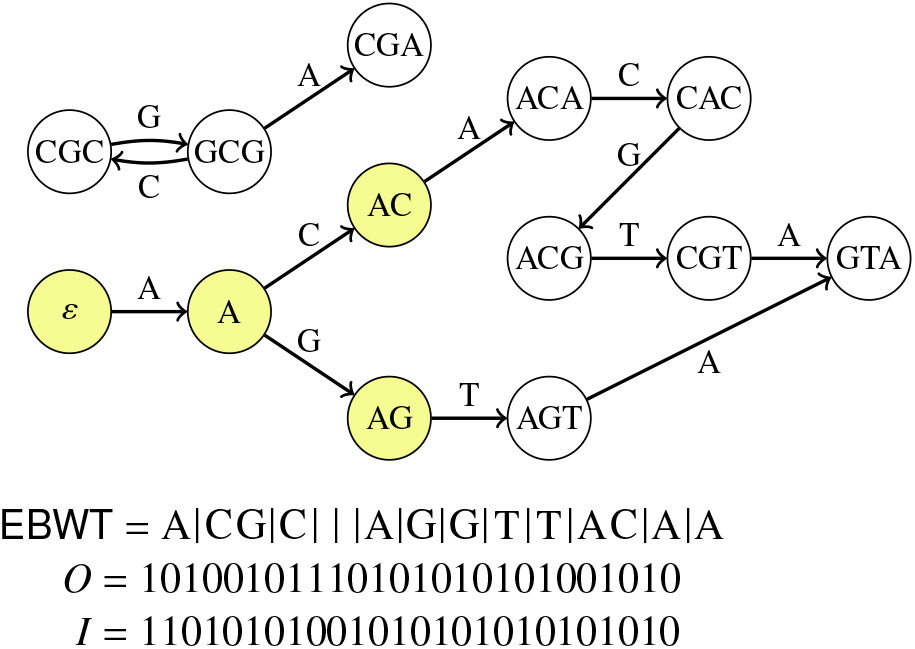
The Wheeler graph on the right corresponds to the edge centric de Bruijn graph with edgemers of length 4 from the set of strings ACGTA, ACACGT, AGTA and GCGCGCGA. The dummy node tree is highlighted in yellow. The colexicographic order of the nodes is the following: ε, A, ACA, CGA, GTA, AC, CAC, CGC, AG, ACG, GCG, AGT, CGT.

The colexicographic ordering implies that if *v* and *u* are nodes such that *v* ≺ *u*, and both have an outgoing edge with the same character label *c* ∈Σ leading to destinations *v′* and *u′*, respectively, then *v′* ≺ *u′*. This means that if we have an interval [*i, j*] of nodes in the order, and follow all outgoing edges from nodes in the interval with the same label *c*, we arrive at another contiguous interval [*i′, j′*] of nodes. This, in turn, implies that it’s enough to compute just the endpoints *I ′* and *j′* because all the other destinations will fall in between them. This can be done in *O*(log σ)-time using rank queries on EBWT and rank/select queries on the indegree and outdegree bit vectors (Gagie et al., 2017). Armed with this, we can locate the interval of nodes at the ends of paths labeled with any pattern *P* in *O*(|*P*| log σ) time by starting from the interval of all nodes, and updating the interval |*P*| times, following the characters of *P*. In the case of the de Bruijn graph, this gives us an algorithm to locate any nodemer in the graph, and to traverse edges in the graph forward and backward.

The original BOSS representation of Bowe et al. (2012) uses the same ideas but the implementation is slightly different. In this representation, nodes with in-degree or outdegree of zero are forbidden. This limitation allows us to represent the information of the indegrees and outdegrees by marking the last outgoing edge from each node and the first incoming edge to each node. This comes at the cost of introducing a new character $ to the alphabet and adding extra edges labeled with $ to EBWT to ensure that every node has at least one outgoing and incoming edge, but it allows a shorter representation of the indegrees and outdegrees.

## 3. Dynamizing Compact Data Structures

BufBOSS follows a line of research dating back to Bentley and Saxe (1980). They proposed an approach to making static data structures semi-dynamic when queries are decomposable, where *semi-dynamic* means we can support additions but not deletions and *decomposable* means we can quickly answer a query about the union of two disjoint sets when given the answers to the queries about those two sets separately. Decomposable queries include membership, minimum, maximum and mean, for example, but not mode.

To support queries and updates to a dynamic set *S* using a static data structure *D*, Bentley and Saxe split *S* into a logarithmic number of disjoint subsets whose sizes are distinct powers of 2 and store an instance of *D* for each subset. To query *S*, we query each instance and combine the results. To insert a new element *x* into *S*, we pool *x* with the elements from the subsets of size 1, 2, 4, etc, destroying the instances of *D* for those subsets as we go, until we find a power of 2 for which there is currently no instance of *D*. We build a new instance of *D* storing the pooled elements, whose number of pooled elements is exactly that power of 2. The dynamic version of *D* for *S* we obtain this way is a constant factor larger than the static version of *D* for *S*, answers queries an *O*(log |*S*|) factor more slowly, and supports each addition in amortized *O*(*P*(*S*) log(|*S*|)/ |*S*|)-time, where *P*(*S*) is the time to build the static version of *D* for *S* .

Overmars and van Leeuwen (1981) extended Bentley and Saxe’s approach to make the amortized complexity of additions worst-case, using copies of the subsets and background processing, and to support deletions. Two techniques for supporting deletions are keeping a “ghost” instance of *D* that holds the deleted elements, and “lazily” deleting elements by marking them in the dynamic version of *D* for *S* and then collapsing a subset and rebuilding when more than half its elements have been lazily deleted.

Munro et al. (2015) adapted Bentley and Saxe’s and Overmars and Van Leeuwen’s results to dynamize compact but static data structures, focusing on indexes for document collections. They proposed keeping a fast but space-inefficient dynamic data structure for the elements that have been added or deleted most recently, as well as a series of compact static data structures for increasingly larger subsets, according to Bentley and Saxe’s scheme; when the dynamic data structure grows too large, they empty it by rebuilding some of the static data structures and incorporating the buffer’s contents. Munro et al. (2015) emphasized that previous approaches to dynamizing compact data structures generally relied on dynamic bitvectors, for which there are fairly strong lower bounds (Fredman and Saks, 1989). By maintaining a dynamic, pointer-based buffer and occasionally rebuilding a compact static data, they can avoid using dynamic bitvectors and thus side-step those lower bounds.

Munro et al. also explained how their ideas could be used to obtain dynamic compact representations of graphs, and Coimbra et al. (2020) recently applied those ideas to obtain an implementation of dynamic *k*^2^-trees that is competitive with previous implementations, which were essentially just static *k*^2^-trees but with dynamic instead of static bitvectors. Since Dynamic-BOSS (Alipanahi et al., 2020a) is essentially the data structure of Bowe et al. (2012) but with dynamic instead of static bit vectors, we are naturally curious how competitive a dynamic de Bruijn graph based on Munro et al.’s ideas will be.

For simplicity and practicality, in this paper we use only one compact static data structure in addition to the dynamic buffer. The main challenge is rebuilding the compact static data structure still in small space, since many data structures that are compact once built are not compact to build, and an implementation is not really compact if it is usually small but balloons every so often.

## 4. Buffering Additions and Deletions

In this section, we describe how to support additions and deletions on a static de Bruijn graph by using a dynamic buffer. Here, we denote *G* = (*V, E*) as the (original) graph before any updates. We denote *A* as the set of edgemers we want to add, and *D* as the set of edgemers we want to delete. The edge set of the modified graph *G′* is *E′* = (*E* ∪*A*) |*D* and the node set *V′* is derived from *E′* as the set of all nodemers that are prefixes or suffixes of edgemers in *E′*.

Since *G* is represented in a BOSS format, the edges and nodes of *G* are identified by their colexicographic ranks. We denote the colexicographic rank of a node *v* ∈*V* with colex(*v*) =|{𝓁 (*u*) ≾ 𝓁 (*v*) : *u* ∈*V*}|, where ≾ denotes the colexicographic comparison and 𝓁 (*v*) is the label of *v*. Similarly, for an edge *e* ∈*E*, we denote colex(*e*) = |{𝓁 (*d*) ≾ 𝓁 (*e*) : *d* ∈*E*}| . We abstract the BOSS structure behind the following interface.

- **Node search:** Given a node label 𝓁 (*v*), return colex(*v*) if *v* ∈ *V* or report that *v* ∉ *V* in *O*(*k* log σ)-time.
- **Edge search:** Given an edge label 𝓁 (*e*), return colex(*e*) if *e* ∈*E* or report that *e* ∉ *E* in *O*(*k* log σ)-time.
- **Out-edge label set:** Given colex(*v*), list the single-character edge labels *L*(*e*) for all edges *e* leaving from *v* in *O*(outdegree(*v*))-time.
- **In-edge label:** Given colex(*v*), return an incoming single-character edge label to *v*, if exists, in *O*(1)-time (by construction, all incoming edges to a node always have the same label).
- **Out-edge rank:** Given colex(*v*) for a node *v* and a character *c*, return the colexicographic rank the edge label 𝓁 (*v*) · *c*, if exists, in *O*(log σ)-time.
- **Forward:** Given the representation of an edge *e* either as its rank colex(*e*), or the pair (colex(*v*), *L*(*e*)), where *v* is the origin of the edge, return colex(*u*), where *u* is the node at the destination of an edge, in *O*(log σ)-time.
- **Backward:** Given colex(*v*) for a node *v*, return colex(*u*), where *u* is a node that has an outgoing edge to *v*, or report that no such edge exists, in *O*(log σ)-time. If there are multiple candidates for *u*, we can return any.
- **In-edge interval:** Given colex(*v*), return the interval [*i, j*] of the colexicographic ranks of the incoming edges to *v* in *O*(1)-time.

See Gagie et al. (2017) for the implementation details of these operations. We now describe how to use these operations and the buffers *A* and *D* to offer graph traversal functionality for *G′*.

We represent the addition buffer *A* with a hash table *H*_*A*_, where the keys are nodemers and the values are the sets of added incoming and outgoing edge labels. If an edge *e* is outgoing from the node then its label is the last character of 𝓁 (*e*); but if it is incoming then its label in the addition buffer is defined as the *first* character of 𝓁 (*e*). We note that the label in the incoming direction is usually not the same as the label of the edge in the Wheeler graph, but rather the label of an incoming edge to a node *u* that is at distance *k* − 2 from the origin of *e* backwards. The keys are packed into integers with log σ bits per character. This allows efficient hashing in *O*(*k* log(σ)/*w*) expected time, where *w* is the width of a machine word in the RAM-machine model. In practice, we limit *k* ≤32, so that with σ = 4, all the keys can be represented with single 64-bit words. The values of *H*_*A*_ are encoded with two bit vectors using σ bits for the forward direction and σ + 1 bits for the backward direction as the special $-symbol is also a possible incoming label.

For example, if we have a buffer nodemer ACA that is contained in buffer edgemers ACAA, ACAG and TACA, then ACA has outgoing labels A and G (the last symbols of the edgemers ACAA and ACAG, where ACA is a prefix) and an incoming label T (the first symbol of the edgemer TACA, where ACA is a suffix). This information is represented in *H*_*A*_[‘ACA’] by bit vectors 00001 in the incoming direction (T is the last character of the incoming alphabet $ACGT) and 1010 in the outgoing direction (A and G are the first and third characters of the outgoing alphabet ACGT). If ACA was not a suffix of any buffer edgemer, then we would have an incoming dollar, which would be encoded by 10000 in the incoming direction

The hash table *H*_*A*_ provides node and edge membership queries to *A* in *O*(*k* log(σ)/*w*) expected time, given a nodemer 𝓁 (*v*) or an edgemer 𝓁 (*e*).

The deletion buffer *D* is represented with just a single bit vector *B*_*D*_ of length |*E*|, such that *B*_*D*_[colex(*e*)] = 1 if and only if *e* ∈*D*. This bit vector provides membership queries to *D* in constant time given colex(*e*), and in *O*(*k* log σ) time given 𝓁 (*e*) by computing colex(*e*) with the BOSS structure.

Adding an edgemer *e* to the addition set *A* is done by splitting *e* into the (*k* − 1)-length prefix and suffix nodemers *v* and *u*, and setting the bits corresponding to the last and first character of *e* in *H*_*A*_[𝓁 (*v*)] and *H*_*A*_[𝓁 (*u*)], respectively. Adding an edgemer *e* to the deletion set *D* is done by searching *e* using the BOSS to compute colex(*e*) and marking the position in *B*_*D*_. We keep the table *H*_*A*_ and the vector *B*_*D*_ synchronized so that when we add an edgemer to *H*_*A*_, we remove it from *B*_*D*_ if present, and vice versa. We do not support node deletions or additions explicitly because the node set is implicitly defined as the set of endpoints of all edges.

We now describe the query interface to the modified graph *G′*. Instead of operating on node identifiers colex(*v*), we operate on node *tokens*, which are pairs (colex(*v*), *H*_*A*_[𝓁 (*v*)]), where the first element of the pair is null if *v* ∉ *V* and the second element is null if 𝓁 (*v*) is not a key of *H*_*A*_.

We can use a token to report whether the represented node exists in *G′*. Given the pair (colex(*v*), *H*_*A*_[𝓁 (*v*)]), the node *v* exists in *G′* if and only if one of the following holds (i) *H*_*A*_[𝓁 (*v*)] is not null (i.e. *v* is a prefix or a suffix of some edge in the addition set), or (ii) colex(*v*) is not null (i.e. *v* is in the BOSS structure) and there is an incoming or outgoing edge to *v* in the BOSS that is not deleted. Checking case (ii) can be implemented by checking for the existence of a 0-bit in *B*_*D*_ in the in-edge range of *v* and at the colexicographic rank of every outgoing edge.

We can list all outgoing edge labels from a node represented by a token (colex(*v*), *H*_*A*_[𝓁 (*v*)]) by taking the union of the BOSS out-edge listing and the edges marked in *H*_*A*_[𝓁 (*v*)], and removing the edges marked in *B*_*D*_.

Forward traversal from a node in *G′* is done by checking the existence of the edge in *G′* using the node token, and then updating colex(*v*) with a BOSS traversal step and *H*_*A*_[𝓁 (*v*)] with hash table lookup.

If the number of dynamic additions and deletions is small, we can optimize the time of graph traversal at the cost of a little space by marking in the BOSS those nodes which are affected by dynamic operations. These are the nodes whose labels are currently keys in *H*_*A*_ or that have at least one incoming or outgoing edge marked in *B*_*D*_. With this, we only have to look up data from *H*_*A*_ and *B*_*D*_ on those nodes, and can rely on the static BOSS most of the time when traversing the graph. The marking can be implemented by using a bit vector *M* such that *M*[*i*] = 1 iff the node with colexicographic rank *i* is marked. This allows us to mark nodes and query whether a node is marked in constant time, given the colexicographic rank of a node. The colexicographic rank of the current node is always known while traversing the graph.

## 5. Batched Updates of Additions and Deletions

In this section, we describe a method to update a BOSS structure with a batch of additions and deletions. The input to the algorithm is a static BOSS structure *G* = (*V, E*), a set *A* of edgemers we want to add, and a set *D* of edgemers we want to delete. The sets *A* and *D* are represented with the hash table *H*_*A*_ and the bit vector *B*_*D*_ described in the previous section. The algorithm requires that *A* ∩*D* = ∅, *A* ∩*E* = ∅ and *D* ⊆*E*, which all hold due to the way *H*_*A*_ and *B*_*D*_ are constructed and synchronized. The output is a BOSS structure containing the set of edgemers of *G* with the set *A* added and *D* deleted.

The update algorithm consists of four phases: (1) a dummy node preparation phase, (2) a merge planning phase, (3) a merge execution phase, which applies both the additions and the deletions, and (4) an optional dummy cleanup phase. We now proceed to describe each phase in this order.

### 5.1. Dummy Node Preparation Phase

In this phase, we iterate over the deletion buffer to check which nodes will be left without an incoming edge after the update. We must add an incoming chain of dummy nodes to these nodes during the update. For every edge that is marked for deletion in *B*_*D*_, we first check whether all incoming edges to its destination node *v* are marked for deletion. If this is the case, we check in the addition buffer whether there are new incoming edges to *v*. If not, we need to add the incoming dummy chain to *v*.

To implement these checks, we need to know *H*_*A*_[𝓁 (*v*)] and the indegree range of *v*. We can find both of these by using the static BOSS structure. We traverse the edge to *v* to find colex(*v*), which allows us compute the indegree range, and then retrieve the label 𝓁 (*v*) by traversing backward *k* − 1 times, which enables us to look up *H*_*A*_[𝓁 (*v*]). All the required new dummy nodes are added to the addition buffer before proceeding to the next phase.

### 5.2. Merge Planning Phase

The purpose of the planning phase is to find the correct places of the node labels in the addition buffer in the colexicographic order of the node labels of the BOSS. Our approach can be seen as a streamlined variant of the planning phase of VariMerge (Muggli et al., 2019), where we have a BOSS structure and a sorted addition buffer rather than two using BOSS structures. This phase does not depend on the deletion buffer at all – the deletions are taken into account in the next phase.

To aid our presentation, we assume that every node label has length *k* −1, by padding the node labels of the dummy nodes with $-symbols from the left. With this, we define the *BOSS matrix*, denoted with *M*_boss_, such that *M*_boss_[*i*][*j*] is the *j*-th character of the *i*-th node label of the BOSS in colexicographic order. Similarly, we define the *buffer matrix*, denoted with *M*_buffer_, such that *M*_buffer_[*i*][*j*] is the *j*-th character of the *i*-th node label of the addition buffer in colexicographic order. We denote with *n*_boss_ and *n*_buffer_ the number of rows in *M*_boss_ and *M*_buffer_ respectively.

We build *M*_buffer_ by extracting all node labels from the addition buffer and sorting them colexicographically. We also attach (as satellite data) to each row the *H*_*A*_ entry encoding the outgoing and incoming labels. However, we note that the matrix *M*_boss_ is not built explicitly. We only define and use it here for explanatory purposes and note that we can use the BOSS structure to access it column by column from right to left. The last column can be constructed by querying the in-edge labels of the nodes in colexicographic order, and Algorithm 2 describes a subroutine that takes as input the *i*-th column of *M*_boss_, and returns the (*i* −1)-th column, by propagating the labels forward in the de Bruijn graph.

After constructing *M*_buffer_, we identify equal rows between *M*_buffer_ and *M*_boss_, and the colexicographic interleaving of the rest of the rows. This is done by running *k* − 1 iterations of a partition refinement subroutine. At the start of iteration *t*, we have a sequence of pairs of half-open intervals ([*a*_1_, *a*_2_), [*b*_1_, *b*_2_)), ([*a*_2_, *a*_3_), [*b*_2_, *b*_3_)), …, ([*a*_*n*−1_, *a*_*n*_), [*b*_*n*−1_, *b*_*n*_)) such that the intervals describe the coarsest partition of rows where every row in the same part has the same suffix of length *t*− 1. That is, the pairs at iteration *t* have the following properties:

1. 1 = *a*_1_ ≤ *a*_2_, …, ≤ *a*_*n*_ = *n*_boss_ + 1 and 1 = *b*_1_ ≤ *b*_2_, …, ≤ *b*_*n*_ = *n*_buffer_ + 1.
2. *M*_boss_[*p*][(*k* − *t* + 1)..(*k* − 1)] = *M*_buffer_[*q*][(*k* − *t* + 1)..(*k* − 1)] for *a*_*i*_ ≤ *p* < *a*_*i*+1_ and *b*_*i*_ ≤ *q* < *b*_*i*+1_.
3. If *a*_*i*_ *≠ a*_*i*+1_, then *M*_boss_[*a*_*i*_][(*k* − *t* + 1)..(*k* − 1)] ≠ *M*_boss_[*a*_*i*+1_][(*k* − *t* + 1)..(*k* − 1)]
4. If *b*_*i*_ *≠ b*_*i*+1_, then *M*_buffer_[*b*_*i*_][(*k* − *t* + 1)..(*k* − 1)] ≠ *M*_buffer_[*b*_*i*+1_][(*k* − *t* + 1)..(*k* − 1)]

To encode these interval pairs succinctly, we only encode the differences *a*_*i*_ − *a*_*i*−1_ and *b*_*i*_ − *b*_*i*−1_ as unary numbers, using in total only *O*(*n*_boss_ + *n*_buffer_) bits. Empty intervals are allowed when a node is present in the BOSS but not in the buffer, or the other way around.

At the start of iteration *t* = 1, we have just a single pair ([1, *n*_boss_ + 1), [1, *n*_buffer_ + 1)). Each iteration refines the partition encoded in the interval pairs by splitting the intervals by the runs of characters in the previous columns of *M*_boss_ and *M*_buffer_. The pseudocode is at Algorithm 3.

#### Algorithm 1

Dummy node preparation phase

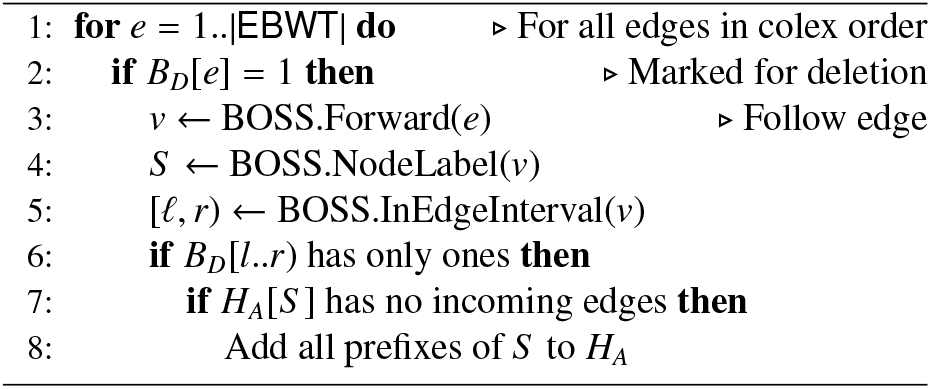

#### Algorithm 2

Subroutine PrevColumn

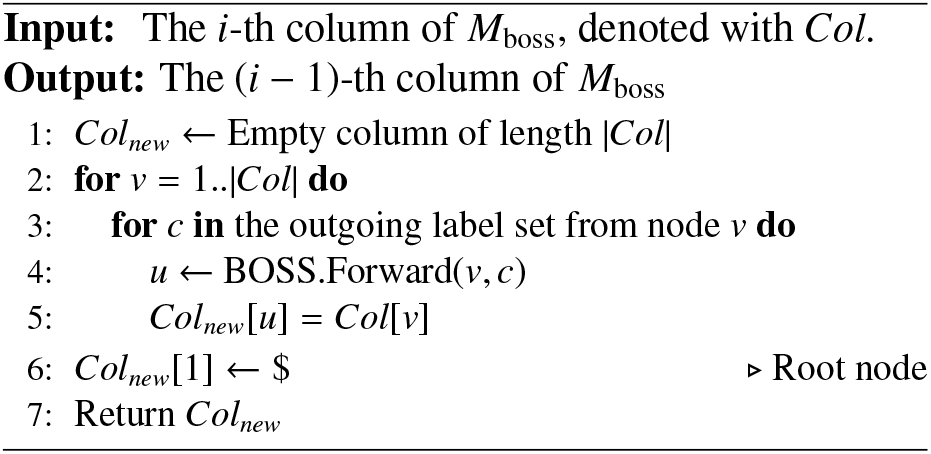

### 5.3. Merge Execution Phase

After the merge planning phase, we move to the execution phase. The interval pairs now identify the nodes that are common to the BOSS and the buffer, and the colexicographic interleaving of all nodes.

In this phase, we stream the interval pairs from the merge planning phase. As we stream the intervals, we enumerate in colexicographic order the indegree and outgoing label set of each node in the buffer and in the BOSS.

During the streaming, the deletion buffer *B*_*D*_ comes into play again. While iterating the nodes, we check things for every node: (1) whether any of the outgoing edges *e* are marked in *B*_*D*_. We can check this by using the BOSS structure to retrieve colex(*e*) and checking *B*_*D*_[colex(*e*)]. (2) Whether any of the incoming edges to the node are marked for deletion: we use the BOSS structure to retrieve the colexicographic range of the incoming edges, and count how many are marked in *B*_*D*_. We remove the outgoing edges that are marked from the outgoing label set, and decrease the indegree by the number of incoming edges that are marked. See Figure 2.

#### Algorithm 3

Merge planning phase

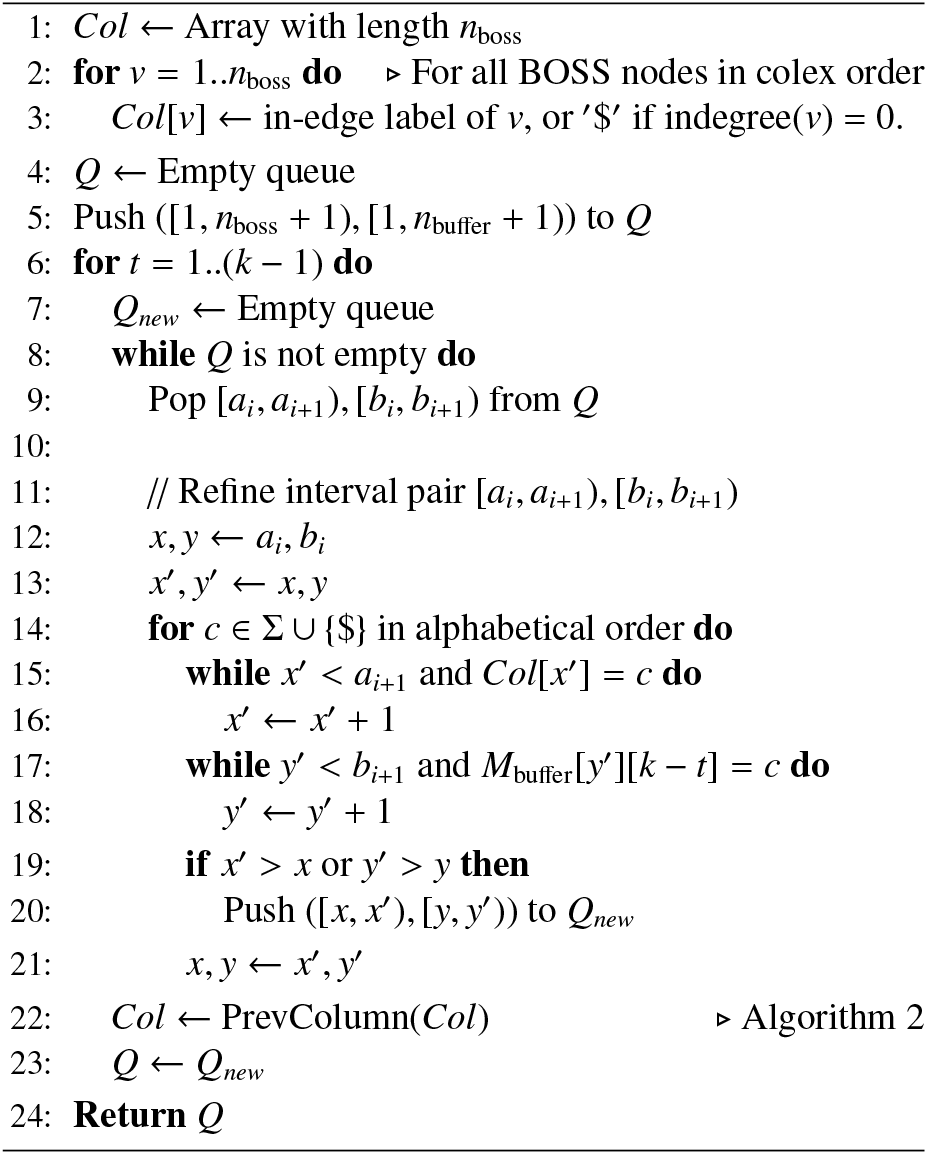

**Figure 2:**
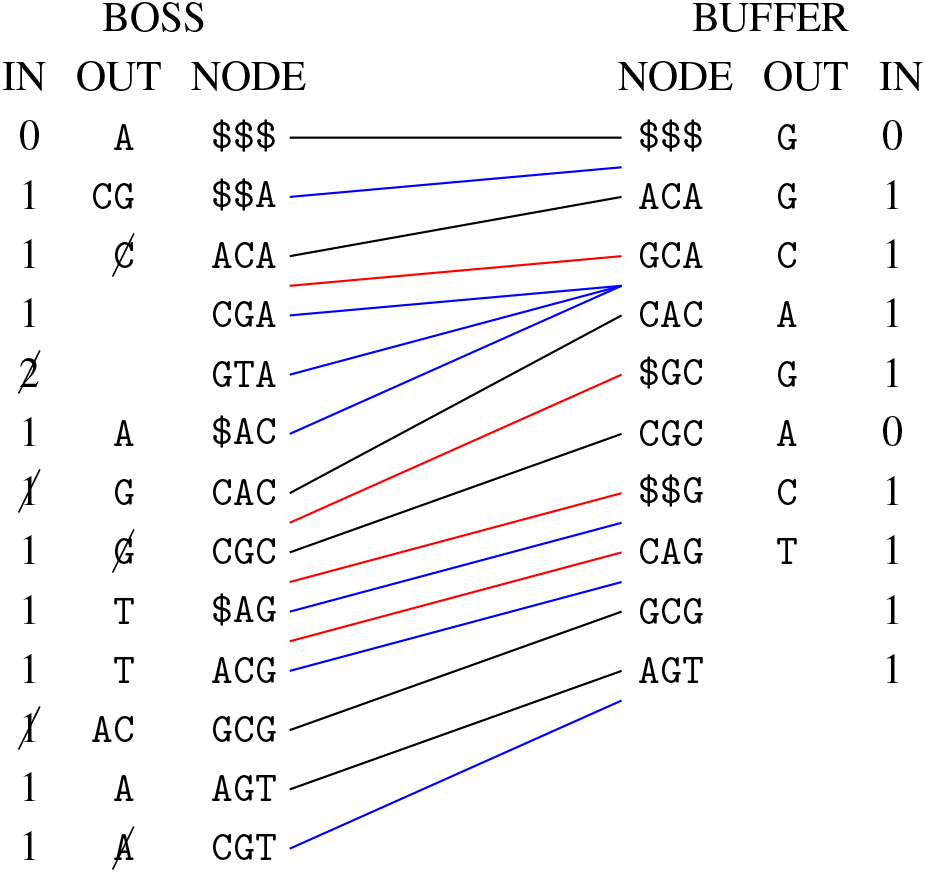
Merge execution phase. In the figure we start from the graph in Figure 1, adding the edgemers in the sequence CGCACAGT and deleting the edgemers CGCG, CGTA and ACAC. The columns IN, OUT and NODE are the indegrees, outgoing edge label sets and the node labels, respectively, including the dummies. The lines in the middle are computed in the planning phase and represented succinctly as a sequence of interval pairs. Black lines connect nodes with the same label. Red lines are for nodes that are present in the buffer but not the BOSS, and blue lines are the other way around. Edges marked for deletion and the indegrees affected by those are marked with a strike-through line.

We stream the new EBWT, *I* and *O* structures of the new updated BOSS to disk while iterating the interval pairs. The pseudocode is at Algorithm 4. The subroutines DeletionsFrom and CountDeletionsTo use the BOSS interface to compute the colexicographic ranks of the incoming and outgoing edges, and access the bit vector *B*_*D*_ to check the deletion flags. The subroutines AdditionsFrom and CountAdditionsTo read from the satellite data attached to the rows of *M*_buffer_ in the planning phase. The subroutine AddNode appends data to the EBWT, *I* and *O* structures of the updated BOSS. Specifically, a call to AddNode(*A, d*), where *A* is an outgoing label set and integer *d* is an indegree, appends 1 · 0^|*A*|^ to the new *O*, 1 · 0^*d*^ to the new *I* and the list *A* to the new EBWT. The structures EBWT, *I* and *O* of the updated BOSS are initialized to empty.

### 5.4. Dummy Cleanup Phase

After the execution phase, we have an optional cleanup phase, where we delete dummy nodes that have become redundant after the addition of the new nodes.

We recall that the dummy nodes form a tree starting from the node of the empty string. We do a depth-first search in the tree of dummies of the updated BOSS from the root using the forward traversal operation in the BOSS interface. When we reach a leaf node of the dummy tree, we follow all the outgoing edges to full nodemers. We check the indegree of each such nodemer. If it is two or more, we can mark the incoming edge from the tree part for deletion, because the target node has another incoming edge which must come from another nodemer. If all out-edges of the dummy node were marked, we can mark the in-edge of the dummy node for deletion as well. Likewise, when we backtrack in the depth-first search, if all out-edges of a dummy tree node are marked for deletion, we mark its in-edge for deletion. This algorithm takes only time proportional to the number of dummy nodes. In the end, we stream the EBWT, *I* and *O* structures of the updated boss, removing marked edges and decrementing the indegrees of the destinations of the deleted edges as done in the execution phase before.

## 6. Results

In this section, we demonstrate the performance of BufBOSS against a variety of dynamic de Bruijn graph implementations from the literature.

### 6.1. Experimental Details

The experiments were run on a server with an Intel Xeon CPU E5-2640 v4 with 40 cores clocked at 2.40GHz, equipped with 755GiB of RAM. The timing and peak memory (RSS) were measured using the Unix utility /usr/bin/time.

### 6.2. Implementation of FDBG-RecSplit

As previously mentioned, we modified FDBG and compare against this modification in addition to original FDBG implementation. Here, we give some background on this modification. Alipanahi et al. (2020a) observed that FDBG fails for datasets approaching 2^32^ distinct nodemers. This is due to hash collisions. FDBG hashes nodemers first with the Karp-Rabin hash function (Karp and Rabin, 1987), and then hashes these hashes with the perfect hash function BBHash (Limasset et al., 2017). The problem is that the Karp-Rabin hash values are stored in 64-bit integers for best compatibility with BBHash, which means that collisions start to happen often at around 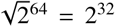 keys due to the birthday paradox phenomenon in probability. In the interest of studying the scalability of FDBG for datasets that have more than 2^32^ nodemers, we edited the implementation to use 128-bit Karp-Rabin hashes, and changed the perfect hashing implementation from BBHash to a recently published new implementation called RecSplit (Esposito et al., 2020a). RecSplit is a good choice for this purpose because it is optimized to work for uniformly random 128-bit keys, and the Karp-Rabin hashes of the nodemers satisfy this model well. Arithmetic with 128-bit values is significantly slower than with 64-bit values, especially when modulo-operations are involved, so we also include results with the original FDBG.

#### Algorithm 4

Merge execution phase

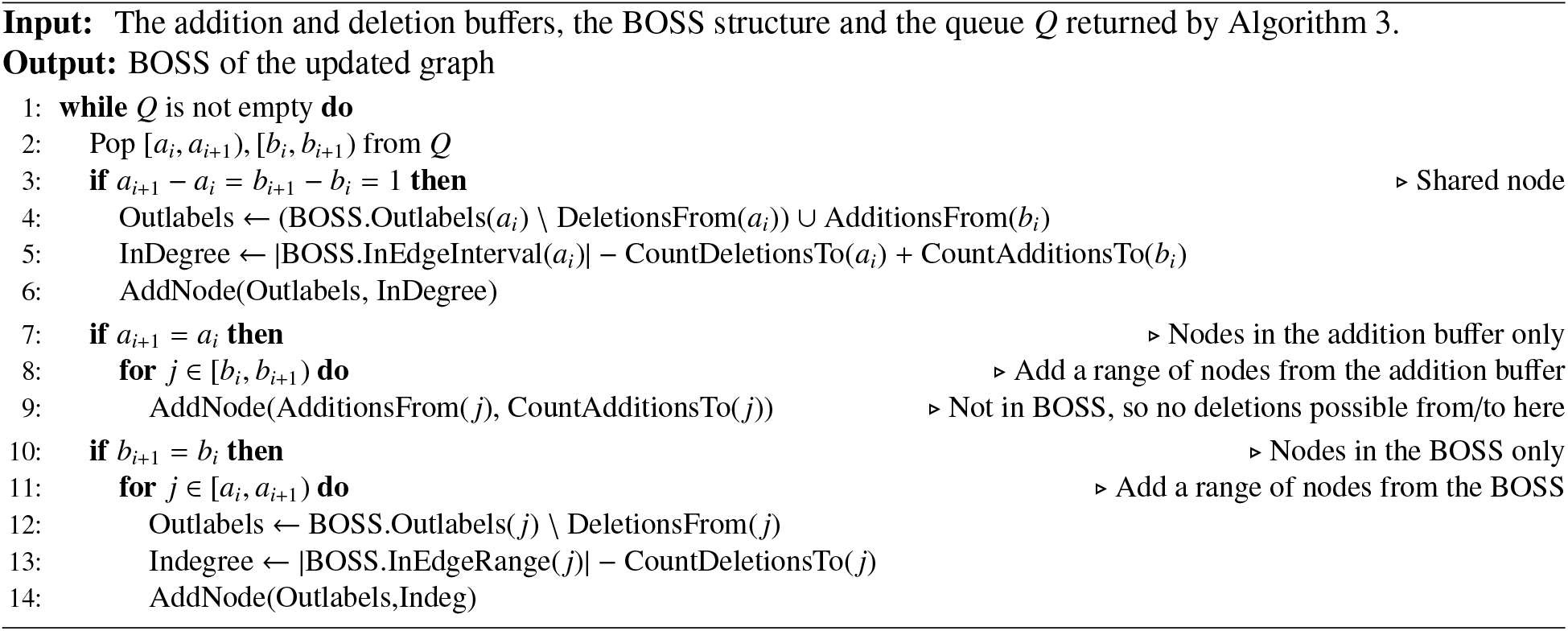

### 6.3. Implementation of BufBOSS

We construct the BOSS structure by first using the KMC3 *k*-mer counter (Kokot et al., 2017) to list all distinct edgemer of the data. KMC3 is highly parallel, using both machine-level parallel vector instructions, and multi-threading to exploit all the cores available on the CPU.

We then write to disk triples (*x, c, b*), where *x* is a nodemer string, *c* is a character either to the left or to the right of *x* in the input data, and *b* is a bit that indicates whether *c* is on the left or on the right. We sort these triples on disk by the colexicographic order of the nodemers *x* using the stxxl library (Dementiev et al., 2008). We then scan the sorted pairs to count the number of characters to the left of *x*. If some nodemer *x* does not have characters on the left, we need to add the dummy nodes corresponding to all prefixes of *x* to the graph. We write these prefixes to another file on disk along with the characters of the right, and sort these colexicographically by the order of the prefixes. Finally, we merge the order of the dummy nodes into the order of the nodemers by streaming the two lists on disk and using logic of the merge phase of merge sort. While we are doing this merge we have, in the correct order, all the information to build the data structure: for every node label *x*, we have the count of characters on the left (the indegree of *x*), and the outgoing edge labels (the outdegree and the labels of the outgoing edges from *x*), all in the colexicographic order of the nodes.

When adding sequences to BufBOSS dynamically, we set a threshold *t* ∈ [0, 1] such that if the addition buffer *H*_*A*_ contains a fraction of more than *t* entries compared to the number of edgemers in the static BOSS structure, we flush the buffers *H*_*A*_ and *B*_*D*_ by running the update algorithm described in Section 5, and clearing the buffers afterwards. This amortizes the cost of individual updates over the buffer flushes, with different time-space trade-offs available for different values of *t*.

The rank and select structures required for queries are implemented with the SDSL-library. The queries are implemented in our Wheeler graph library. The dynamic buffer hash table *H*_*A*_ is implemented with the C++ standard library. Like FDBG, we store edgemers in 64-bit integers, so the maximum allowed edgemer length is 32.

### 6.4. Competing Tools

We compare the performance of BufBOSS to the following dynamic de Bruijn graph implementations: DynamicBOSS (Alipanahi et al., 2020a), FDBG (Crawford et al., 2018), FDBG-RecSplit, and Bifrost (Holley and Melsted, 2020). An overview of recent de Bruijn graph implementations and their attributes is given in Table 1. Hence, we did not compare against BFT (Holley et al., 2015), Vari (Muggli et al., 2017), Rainbowfish (Almodaresi et al., 2017), and Pufferfish (Almodaresi et al., 2018), and Mantis (Pandey et al., 2018) because they cannot perform addition or deletion of data.

We deviate from our experimental setup in one detail to fairly evaluate the time for addition for Bifrost: we discount the time to load the index into memory. This is because Bifrost does not serialize its index to disk but rather writes all maximal non-branching paths of the de Bruijn graph (unitigs) to disk in a graph format. When Bifrost loads the index, it must re-index the whole dataset by computing and hashing *k*-mer minimizers for all unitigs in the data.

### 6.5. Datasets

Our first set of datasets is short reads from the bacteria E. coli. We used 28 428 648 paired-end reads generated from whole genome sequencing of E.coli K-12 substr. MG1655 dataset (NCBI SRA accession ERX002508). We refer to this as 28M-e. Next, we split this dataset to generate smaller datasets of sizes 2 000, 200 000, 2 000 000, and 14 000 000 reads, which we denote by 20K-e, 200K-e, 2M-e and 14M-e, respectively.

The second set of inputs is a series of metagenomic read sets of increasing size from a study on antimicrobial resistant determinants in commercial beef production (Noyes et al., 2016). This dataset consists of 87 datasets that were generated by sequencing DNA that was collected from various locations across a beef production cycle – the goal being to identify specific points where the pathogenic load either increased or decreased and thus, determine of the effectiveness of the existing interventions used to reduce pathogenic load. The NCBI SRA accession number for these datasets is PRJNA292471. From these 87 datasets, we selected the datasets with the smallest and largest number of reads, which 55 242 004 and 11 136 890 sequence reads, respectively. We refer to as 55M and 11M. We generated smaller datasets by randomly selecting 20 000, 200 000 and 2 000 000 reads from 11M dataset which we refer to these as 20K, 200K and 2M, respectively. Next, we concatenated three datasets that consist of 55 242 004, 44 035 852, and 52 833 978 reads to create a dataset with over 150 million reads, which we refer to as 150M. Lastly, we added 7 different datasets to 150M to generate a dataset with over 600 million reads, which we refer to as 600M.

Since most of the tools we experiment with do not understand characters outside of the DNA-alphabet {A,C,G,T}, such as the invalid character N, we split the reads into pieces that contain only characters from the DNA alphabet. Since FDBG, FDBG-Recsplit and DynamicBOSS do not index reverse complements, but BufBOSS and Bifrost do, we concatenate the input read sets with their reverse complements for the former three tools to make the graphs the same for all tools.

### 6.6. Construction

For each E. coli dataset, we build the index of each tool using 50% of the reads, then add the next 25% of the reads and finally delete the remaining 25%. Addition and deletion are shown in the next subsection. Here, we discuss the construction time, space and memory. Table 3 shows the time and peak memory for construction as well as the size of the final data structure on disk. In addition, Figures 3a and 3b illustrate the time versus memory required for construction of the data structure using the largest E. coli dataset, and the time versus disk space required for construction of the data structure using the largest E. coli dataset.

**Figure 3:**
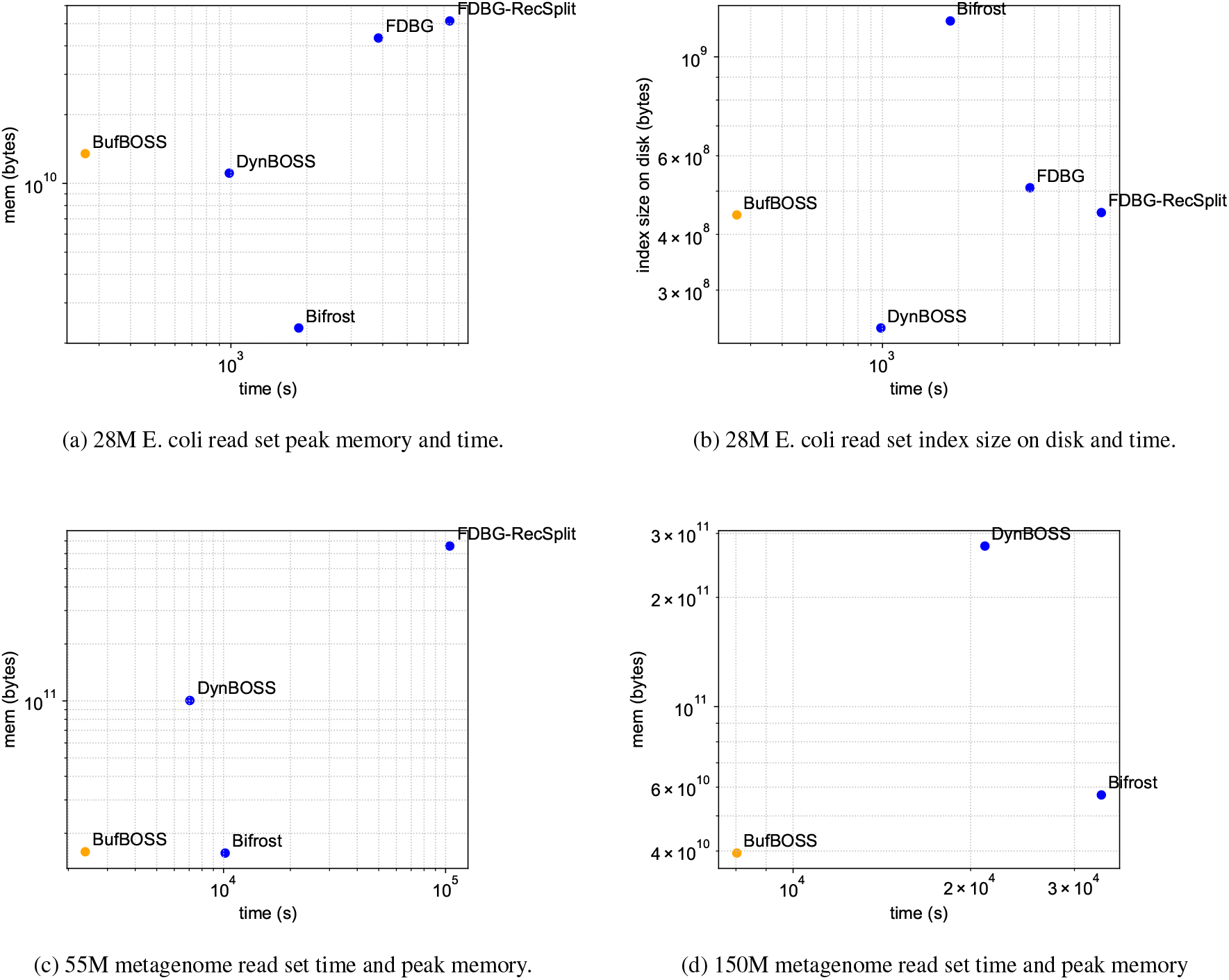
Index construction.

The *k*-mer preprocessing steps of all tools are included in the time and peak memory numbers. BufBOSS was consistently the fastest index to construct, being almost 4 times faster than the closest competitor DynamicBOSS on the largest dataset. Our index construction takes a lot of memory on the small datasets, but this is due to KMC3 always requiring a large amount of memory even on small datasets. When the size of the data increases, our peak memory becomes more competitive. Bifrost had the smallest memory by a factor of 5 to the closest competitors DynamicBOSS and BufBOSS. DynamicBOSS had the smallest index on disk, being half the size of BufBOSS. This is due to the BOSS-implementation of DynamicBOSS being geared more toward small size rather than speed.

We test further scalability of construction using the metagenome datasets, omitting the original FDBG as it can not handle more than 2^32^ nodemers. The data for the rest of the tools is in Table 2. Figure 3c shows the time-memory plot for the 55M dataset, which is the largest FDBG-Recsplit was able to process before running out of memory, and Figure 3d shows the same plot for the 150M dataset without FDBG-Recsplit.

**Table 2:** Index construction on the metagenome datasets. The units of disk and peak memory are GB (2^30^ bytes) and the format of the time is hours:minutes:seconds. FDBG is not included because it runs into hash collisions for large datasets. FDBG-RecSplit ran out of memory (OOM) on the 150M dataset. The edgemer packing preprocessing step of DynamicBOSS ran out of memory on the 600M dataset. The number of distinct canonical edgemers in these datasets in increasing order of size are 1 203 852, 11 839 506, 107 122 222, 478 210 723, 1 360 576 988, 3 741 452 498 and 11 676 829 812. (c) 55M metagenome read set time and peak memory. (d) 150M metagenome read set time and peak memory

|  | BufBOSS |  |  | DynamicBOSS |  |  | FDBG-RecSplit |  |  | Bifrost |  |  |
| --- | --- | --- | --- | --- | --- | --- | --- | --- | --- | --- | --- | --- |
|  | Disk | Memory | Time | Disk | Memory | Time | Disk | Memory | Time | Disk | Memory | Time |
| 20K | 0.00 | 5.26 | 00:00:08 | 0.00 | 0.37 | 00:00:30 | 0.00 | 0.52 | 00:00:41 | 0.00 | 0.02 | 00:00:07 |
| 200K | 0.03 | 6.50 | 00:00:23 | 0.02 | 2.45 | 00:01:44 | 0.04 | 4.77 | 00:07:03 | 0.02 | 0.10 | 00:01:08 |
| 2M | 0.30 | 10.07 | 00:02:58 | 0.17 | 7.64 | 00:08:26 | 0.38 | 42.04 | 01:15:17 | 0.19 | 1.42 | 00:10:22 |
| 11M | 1.32 | 12.82 | 00:13:24 | 0.73 | 33.54 | 00:35:11 | 1.68 | 183.74 | 07:24:03 | 1.13 | 6.00 | 00:53:12 |
| 55M | 3.65 | 14.88 | 00:39:42 | 2.01 | 93.66 | 01:57:32 | 4.69 | 612.64 | 29:00:39 | 4.76 | 14.64 | 02:49:29 |
| 150M | 10.00 | 36.77 | 02:13:47 | 5.52 | 258.03 | 05:52:31 | OOM | OOM | OOM | 14.39 | 53.18 | 09:15:26 |
| 600M | 30.67 | 112.55 | 09:18:19 | OOM | OOM | OOM | OOM | OOM | OOM | 59.08 | 162.00 | 45:30:00 |

**Table 3:** Construction on the E. coli dataset. The units of disk and peak memory are GB (2^30^ bytes) and the format of the time is hours:minutes:seconds. The number of distinct canonical edgemers in these datasets in increasing order of size are 626 875, 4 087 049, 13 241 253, 63 704 311 and 128 431 292.

|  | BufBOSS |  |  | DynamicBOSS |  |  | FDBG-RecSplit |  |  | FDBG |  |  | Bifrost |  |  |
| --- | --- | --- | --- | --- | --- | --- | --- | --- | --- | --- | --- | --- | --- | --- | --- |
|  | Disk | Memory | Time | Disk | Memory | Time | Disk | Memory | Time | Disk | Memory | Time | Disk | Memory | Time |
| 20K-e | 0.00 | 5.15 | 00:00:07 | 0.00 | 0.11 | 00:00:44 | 0.00 | 0.27 | 00:00:21 | 0.00 | 0.23 | 00:00:07 | 0.00 | 0.02 | 00:00:04 |
| 200K-e | 0.01 | 5.78 | 00:00:12 | 0.01 | 1.12 | 00:00:54 | 0.01 | 1.54 | 00:02:27 | 0.02 | 1.30 | 00:00:51 | 0.01 | 0.05 | 00:00:25 |
| 2M-e | 0.04 | 6.97 | 00:00:25 | 0.02 | 2.61 | 00:02:16 | 0.04 | 5.14 | 00:09:37 | 0.05 | 4.36 | 00:04:20 | 0.08 | 0.16 | 00:02:01 |
| 14M-e | 0.20 | 10.07 | 00:02:06 | 0.11 | 5.10 | 00:09:18 | 0.21 | 23.74 | 00:54:19 | 0.24 | 19.98 | 00:29:09 | 0.54 | 1.09 | 00:13:36 |
| 28M-e | 0.41 | 12.57 | 00:04:24 | 0.23 | 10.33 | 00:16:25 | 0.42 | 47.87 | 02:02:57 | 0.47 | 40.30 | 01:04:05 | 1.12 | 2.17 | 00:30:57 |

The external memory construction of BufBOSS now starts to show its scalability on the largest dataset, with the lowest peak memory out of all tools, while maintaining the fastest construction. DynamicBOSS again has the smallest index size on disk. The peak construction memory of FDBG-Recsplit is very large, and exceeds the 755GiB memory capacity of the machine already on the 150M dataset. This enormous peak memory is probably caused by the fact that FDBG-Recsplit holds all input edgemers in memory while building the index. The *k*-mer preprocessing step of dynamicBOSS also exceeded the 755GiB capacity of the machine on the 600M dataset.

### 6.7. Addition and Deletion

Since only two tools out of five are able to scale on construction of the metagenome datasets, we benchmark additions, deletions and queries only on the E. coli dataset. For the E. coli datasets, we evaluated the resources needed to add an additional 25% of the reads, and delete the remaining 25% of the reads. We added and deleted using the graphs that were constructed for the previous section. For deletion, we only compare to FDBG-Recsplit, FDBG and DynamicBOSS since Bifrost does not perform deletion. Tables 4 and 5 illustrate the time, memory and disk required for addition and deletion, respectively. Also, Figures 4 and 5 illustrate the time versus space for additions and deletions, respectively.

**Table 4:** Additions after construction of 3 on the E. coli dataset. The units of disk and peak memory are GB (2^30^ bytes) and the format of the time is hours:minutes:seconds. BufBOSS uses the buffer fraction parameter 0.025. The number of distinct canonical edgemers in these datasets in the addition set in increasing order of size are 329 523, 2 501 881, 9 047 060, 38 892 710 and 56 315 888.

|  | BufBOSS |  |  | DynamicBOSS |  |  | FDBG-RecSplit |  |  | FDBG |  |  | Bifrost |  |  |
| --- | --- | --- | --- | --- | --- | --- | --- | --- | --- | --- | --- | --- | --- | --- | --- |
|  | Disk | Memory | Time | Disk | Memory | Time | Disk | Memory | Time | Disk | Memory | Time | Disk | Memory | Time |
| 20K-e | 0.00 | 0.02 | 00:00:43 | 0.00 | 0.09 | 00:12:51 | 0.01 | 0.32 | 00:00:56 | 0.01 | 0.32 | 00:00:21 | 0.00 | 0.03 | 00:00:03 |
| 200K-e | 0.01 | 0.09 | 00:02:47 | 0.01 | 0.64 | 00:59:22 | 0.05 | 1.42 | 00:08:57 | 0.05 | 1.42 | 00:03:13 | 0.01 | 0.07 | 00:00:18 |
| 2M-e | 0.06 | 0.29 | 00:13:43 | 0.03 | 2.47 | 04:22:04 | 0.20 | 9.89 | 01:31:27 | 0.20 | 9.89 | 00:30:02 | 0.12 | 0.38 | 00:01:21 |
| 14M-e | 0.33 | 1.33 | 01:37:50 | 0.16 | 11.42 | 27:08:14 | 1.26 | 67.13 | 11:38:54 | 1.29 | 67.16 | 04:04:59 | 0.81 | 1.56 | 00:08:08 |
| 28M-e | 0.56 | 2.26 | 02:28:24 | 0.30 | 17.79 | 40:33:54 | 1.88 | 120.07 | 23:08:40 | 1.94 | 120.13 | 07:29:05 | 1.54 | 2.63 | 00:14:37 |

**Table 5:** Deletions after construction of table 3 and additions of table 4 on the E. coli dataset. The units of disk and peak memory are GB (2^30^ bytes) and the format of the time is hours:minutes:seconds. Bifrost is not included because it does not support deletions. The number of distinct canonical edgemers in these datasets in the deletion set in increasing order of size are 330 563, 2 526 285, 8 786 286, 4 3705 141 and 48 377 370.

|  | BufBOSS |  |  | DynamicBOSS |  |  | FDBG-RecSplit |  |  | FDBG |  |  |
| --- | --- | --- | --- | --- | --- | --- | --- | --- | --- | --- | --- | --- |
|  | Disk | Memory | Time | Disk | Memory | Time | Disk | Memory | Time | Disk | Memory | Time |
| 20K-e | 0.00 | 0.01 | 00:00:01 | 0.00 | 0.09 | 00:03:32 | 0.01 | 0.22 | 00:00:50 | 0.01 | 0.22 | 00:00:17 |
| 200K-e | 0.01 | 0.04 | 00:00:07 | 0.01 | 0.64 | 01:44:39 | 0.10 | 1.22 | 00:28:53 | 0.10 | 1.22 | 00:12:50 |
| 2M-e | 0.06 | 0.13 | 00:01:30 | 0.03 | 2.42 | 04:52:23 | 0.34 | 7.84 | 01:28:27 | 0.34 | 7.85 | 00:39:22 |
| 14M-e | 0.33 | 0.79 | 00:17:24 | 0.18 | 12.62 | 12:46:43 | 1.56 | 51.69 | 05:49:05 | 1.59 | 51.72 | 03:23:58 |
| 28M-e | 0.56 | 1.23 | 00:32:32 | 0.32 | 15.96 | 17:23:39 | 2.31 | 94.22 | 14:35:19 | 2.37 | 94.28 | 08:56:51 |

**Figure 4:**
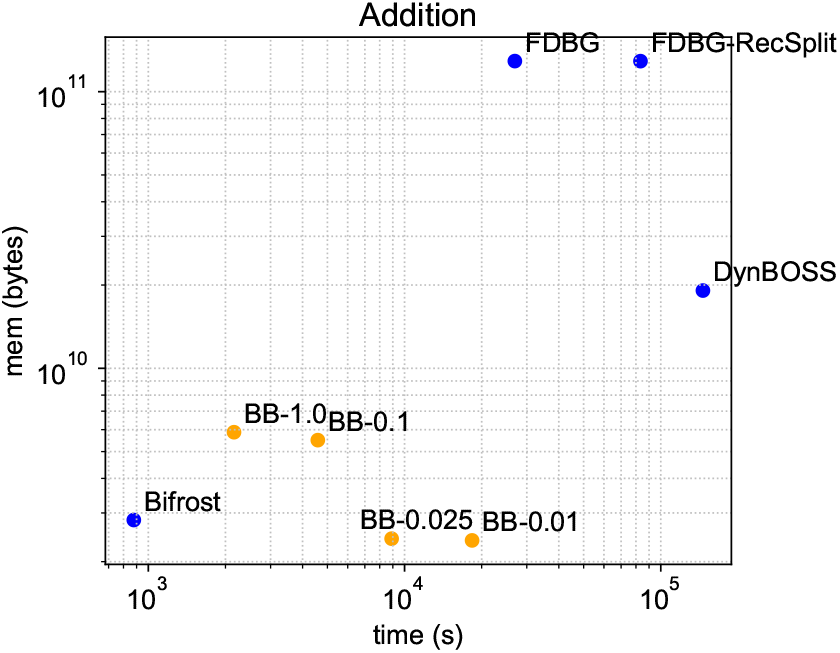
Addition performance on the 28M E. coli dataset. The data points labeled with BB-*t* are runs on BufBOSS with different buffer fraction parameters *t*.

**Figure 5:**
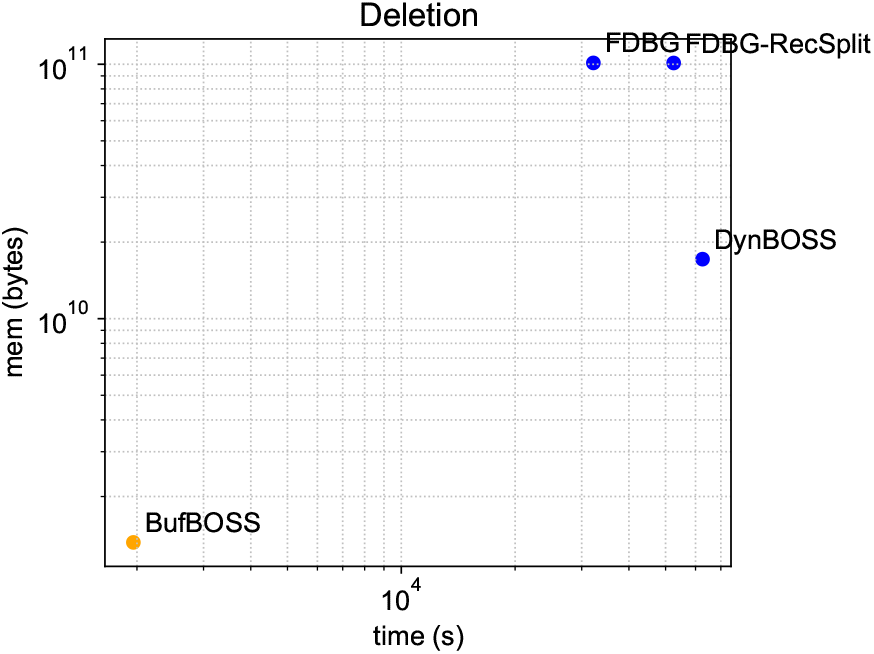
Deletion performance on the 28M E. coli dataset. Here we do not vary the buffer fraction parameter for BufBOSS because buffer flushes are not needed when doing only deletions.

As previously mentioned, we discount the time to load the index into memory for Bifrost. This loading takes two minutes on the 28M E. coli read set. The loading time of other tools is negligible in our experiment. Hence, we subtract Bifrost’s loading time out to avoid skewing the addition performance results.

Figure 4 shows a few different time-space trade-offs BufBOSS can achieve by varying the buffer flushing threshold *t*. With the values of *t* = 0.025 or *t* = 0.01, BufBOSS achieves the lowest peak memory out of all tools, losing in time only to Bifrost.

FDBG and FDBG-Recsplit perform very poorly with additions, because they store new nodemers in a hash table as strings. This blows up both the peak memory and the size of the index on disk. The smallest index on disk is DynamicBOSS, being about half as small as BufBOSS in the end. Bifrost has the fastest additions even when BufBOSS does not flush the buffer at all (*t* = 1.0).

In deletion efficiency, BufBOSS is by far superior, being an order of magnitude better than the other tools in both time and peak space. The deletions of BufBOSS are very efficient because we only mark the deleted edgemers in a bit vector. No buffer flushes are required since the space of the deletion buffer bit vector *B*_*D*_ does not depend on the number of deletions. We note that the other tools could adopt a similar technique for deletions.

### 6.8. Edge Existence Queries

We test the existence query speeds of the tools on six different types of input data. The first three query types are single edgemers: we query edgemers sampled from the construction read set, the addition read set, and completely randomly generated edgemers which do not exist in the data with high probability. The last three are the same, except that instead of single edgemers, we query all edgemers of whole read sequences in the same query, which speeds up the query time per edgemer in BufBOSS, because if the queries exist in the data, we can just traverse the de Bruijn graph instead of searching all the edgemers separately.

All the queries are done on the index with the construction and addition edgemers, but without having deleted the final 25% of edgemers, because for consistency, we want all the queries to be done on the same de Bruijn graph, and Bifrost can not delete edgemers.

Again we factor out the time to load the index for all tools. Figure 6 shows the average query times on the different types of queries. Bifrost has the fastest queries on all query types, with BufBOSS coming in second. DynamicBOSS queries are significantly slower than those of BufBOSS because of the heavy price DynamicBOSS pays for using dynamic bit vectors in the implementation. We see that BufBOSS benefits significantly if the queries are given as whole reads rather than single edgemers, because if the reads are in the graph, we do not have to search every edgemer from scratch, but can traverse the graph instead. This improves the time complexity of querying all edgemers of a read of length *m* from *O*(*km* log σ) to *O*(*m* log σ). Querying edgemers from the set of added edgemers did not make a noticeable difference compared to querying from the initial construction set. Random edgemers are faster to query than single existing edgemers, because the search can terminate early when a prefix of the edgemer is found to not exist in the graph, yet still slower than querying whole existing reads because of the efficiency of graph traversal.

**Figure 6:**
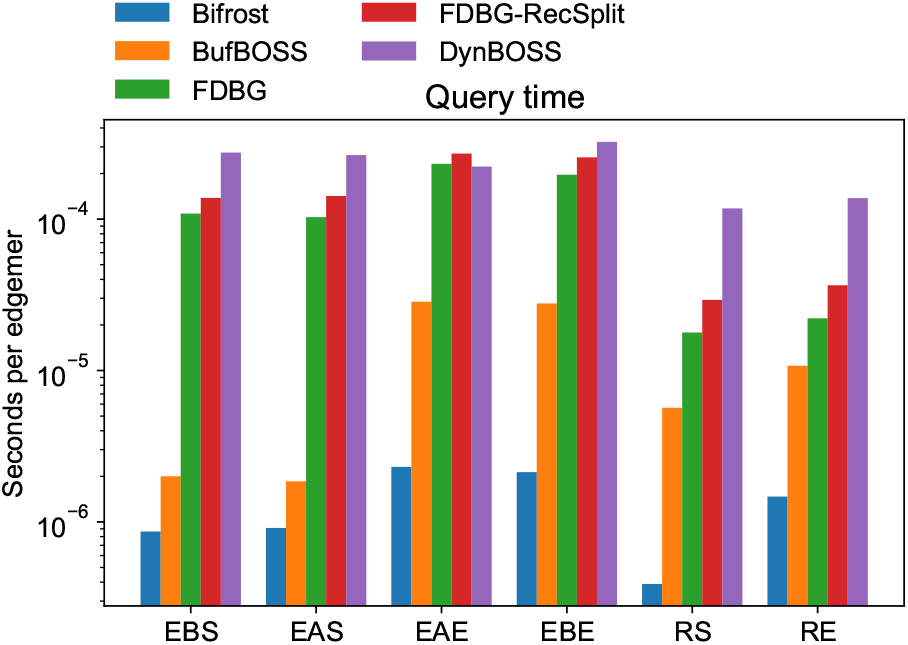
Edge existence queries for six different types of edgemers against the 28M E. coli index including the added sequences. The query types are the following: EBS = existing build sequence, EAS = existing added sequence, EAE = existing added edgemer, EBE = existing build edgemer, RS = random sequence, RE = random edgemer.

The query efficiency of Bifrost is explained partly by good memory locality. Bifrost finds the unitig containing the current edgemer, and can scan forward sequentially in memory in the unitig. On the other hand, the BOSS structure has to jump around in memory in an unpredictable way, resulting in a large number of cache misses. FDBG and FDBG-Recsplit face a similar issue because adjacent edgemers being stored by their hash values in unpredicatable memory locations. Sirén et al. (2020) recently developed a technique to improve BWT-based data structures’ memory locality, but so far it has been applied only in the context of variation graphs.

## 7. Discussion and Future Work

We have shown that buffering updates into a BOSS data structure can provide attractive trade-offs in terms of time, memory and disk usage compared to other tools. In particular, our approach to deletion is clearly the most efficient method for deletion in a de Bruijn graph. Our index construction was also the fastest, which is mainly due to the efficiency of the libraries KMC3 and stxxl used for edgemer listing and sorting. An important caveat in construction is that while BufBOSS was ran single-threaded as the other tools, the external libraries KMC3 and stxxl used for BufBOSS construction use parallelism nonetheless.

Observing the source code of the tools reveals short-comings in the implementations of all the tools. For example, the Bifrost index on disk is just a representation of the de Bruijn graph as a collection of unitigs and edge pointers, but it does not include a indexing structure and thus, Bifrost has to rebuild the index every time it is run. FDBG, FDBG-Recsplit and DynamicBOSS load all the distinct edgemers into memory at once for additions and deletions, even though this should not be needed. All the tools could in principle implement deletions efficiently by simply marking the deleted edgemers, like BufBOSS does. These shortcomings indicate that much could still be gained by more careful implementations of the methods, and that the results we obtained do not necessarily reflect the fundamental limitations of the approaches. Bifrost is the most mature implementation as the codebase dates back to 2011.

The implementation of BufBOSS could also be improved. Recently, Egidi et al. (2020) consider Wheeler graph merging in addition to de Bruijn graph merging, and although, they show that the problem is computationally more challenging, they propose an algorithm specific to BOSS that has a lower peak space than VariMerge in theory. It is worth noting that these results are still only theoretical and they do not provide any implementation. Nonetheless, the practicality of their results warrants future investigation.

More drastic modifications possible to BufBOSS include applying the technique of Sirén et al. (2020) for improving memory locality, and maintaining a dynamic longest-common prefix (LCP) array to allow us to change the order *k* of the graph while navigating it, up to some maximum order set at construction time. Previously, Boucher et al. (2015) showed how the BOSS data structure can be augmented to support variable orders; Belazzougui et al. (2016b) and Belazzougui et al. (2018) showed how the resulting data structure can be made bidirectional; and Díaz-Domínguez et al. (2019) showed how the LCP array can be replaced by a succinct tree shape, at the cost of the order no longer being known.

Although there are tools that recommend the best fixed order for a de Bruijn graph (Chikhi and Medvedev, 2014), variations in read coverage, particularly from single-cell sequencing, can mean that no single order is appropriate for an entire genome or set of genomes. Some tools, such as SPAdes (Bankevich et al., 2012) and IDBA (Peng et al., 2010), use several iterations with different orders, but rebuilding massive graphs this way would be impractical and defeat the purpose of the dynamization. As far as we know, there is currently no algorithm to maintain a dynamic LCP array, it seems like a reasonable extension of BufBOSS. In contrast, it seems unlikely that Bifrost, for example, can be made to support variable orders without greatly increasing its memory usage.

## References

Alipanahi, B., Kuhnle, A., Puglisi, S., Salmela, L., Boucher, C., 2020a. Succinct Dynamic de Bruijn Graphs. Bioinformatics btaa546.

Alipanahi, B., Muggli, M., Jundi, M., Noyes, N., Boucher, C., 2020b. Metagenome SNP calling via read-colored de Bruijn graphs. Bioinformatics btaa081.

Alipanahi, B., Salmela, L., Puglisi, S.J., Muggli, M., Boucher, C., 2017. Disentangled long-read de Bruijn graphs via optical maps, in: Proc of WABI, pp. 1:1–1:14.

Allard, M.W., Strain, E., Melka, D., Bunning, K., Musser, S.M., Brown, E.W., Timme, R., 2016. Practical value of food pathogen traceability through building a whole-genome sequencing network and database. J Clin Microbiol 54, 1975–1983.

Almodaresi, F., Pandey, P., Patro, R., 2017. Rainbowfish: A succinct colored de Bruijn graph representation, in: Proc of WABI, pp. 251– 265.

Almodaresi, F., Sarkar, H., Srivastava, A., Patro, R., 2018. A space and time-efficient index for the compacted colored de Bruijn graph. Bioinformatics 34, i169–i177.

Bankevich, A., Nurk, S., Antipov, D., Gurevich, A.A., Dvorkin, M., Kulikov, A.S., Lesin, V.M., Nikolenko, S.I., Pham, S., Prjibelski, A.D., et al., 2012. SPAdes: a new genome assembly algorithm and its applications to single-cell sequencing. J Comput Biol 19, 455–477.

Belazzougui, D., Gagie, T., Mäkinen, V., Previtali, M., 2016a. Fully Dynamic de Bruijn Graphs, in: Proc of SPIRE, pp. 145–152.

Belazzougui, D., Gagie, T., Mäkinen, V., Previtali, M., Puglisi, S.J., 2016b. Bidirectional variable-order de Bruijn graphs, in: Proc of LATIN, Springer. pp. 164–178.

Belazzougui, D., Gagie, T., Mäkinen, V., Previtali, M., Puglisi, S.J., 2018. Bidirectional variable-order de bruijn graphs. Int J Found Comput Sci 29, 1279–1295.

Bentley, J.L., Saxe, J.B., 1980. Decomposable searching problems I: Static-to-dynamic transformation. J Algo 1, 301–358.

Boucher, C., Bowe, A., Gagie, T., Puglisi, S.J., Sadakane, K., 2015. Variable-order de Bruijn graphs, in: Proc of DCC, pp. 383–392.

Bowe, A., Onodera, T., Sadakane, K., Shibuya, T., 2012. Succinct de Bruijn graphs, in: Proc of WABI, pp. 225–235.

Cameron, D., Schröder, J., Sietsma Penington, J., Do, H., Molania, R., Dobrovic, A. Speed, T., Papenfuss, A., 2017. GRIDSS: sensitive and specific genomic rearrangement detection using positional de Bruijn graph assembly. Genome Res 27, 2050–2060.

Chikhi, R., Limasset, A., Medvedev, P., 2016. Compacting de Bruijn graphs from sequencing data quickly and in low memory. Bioinformatics 32, i201–i208.

Chikhi, R., Medvedev, P., 2014. Informed and automated k-mer size selection for genome assembly. Bioinformatics 30, 31–37.

Coimbra, M.E., Francisco, A.P., Russo, L.M., De Bernardo, G., Ladra, S., Navarro, G., 2020. On dynamic succinct graph representations, in: Proc of DCC, pp. 213–222.

Crawford, V., Kuhnle, A., Boucher, C., Chikhi, R., Gagie, T., 2018. Practical Dynamic de Bruijn Graphs. Bioinformatics 34, 4189– 4195.

Danko, D., et al., 2021. Global genetic cartography of urban metagenomes and anti-microbial resistance. Cell 184, 1–18.

Dementiev, R., Kettner, L., Sanders, P., 2008. STXXL: standard template library for xxl data sets. Softw Pract Exp 38, 589–637.

Díaz-Domínguez, D., Gagie, T., Navarro, G., 2019. Simulating the DNA overlap graph in succinct space, in: Proc of CPM, pp. 26:1– 26:20.

Egidi, L., Louza, F., Manzini, G., 2020. Space efficient merging of de Bruijn graphs and wheeler graphs. arXiv .

Esposito, E., Graf, T.M., Vigna, S., 2020a. RecSplit: Minimal perfect hashing via recursive splitting, in: Proc of ALENEX, pp. 175–185.

Esposito, E., Mueller-Graf, T., Vigna, S., 2020b. RecSplit: Minimal Perfect Hashing via Recursive Splitting, in: Proc of ALENEX, pp.175–185. doi:10.1137/1.9781611976007.14.

Ferragina, P., Manzini, G., 2005. Indexing compressed text. JACM 52, 552–581.

Fredman, M., Saks, M., 1989. The cell probe complexity of dynamic data structures, in: Proc of STOC, pp. 345–354.

Gagie, T., Manzini, G., Sirén, J., 2017. Wheeler graphs: A framework for BWT-based data structures. Theor Comput Sci 698, 67–78.

Holley, G., 2019. Personal email communication with authors of BFT.

Holley, G., Melsted, P., 2020. Bifrost–highly parallel construction and indexing of colored and compacted de Bruijn graphs. Genome Bio 21, 249.

Holley, G., Wittler, R., Stoye, J., 2015. Bloom filter trie–a data structure for pan-genome storage, in: Proc. of WABI, pp. 217–230.

Iqbal, Z., Caccamo, M., Turner, I., Flicek, P., McVean, G., 2012. De novo assembly and genotyping of variants using colored de Bruijn graphs. Nat Genet 44, 226–232.

Karp, R.M., Rabin, M.O., 1987. Efficient randomized pattern-matching algorithms. IBM J Res Dev 31, 249–260.

Kokot, M., Długosz, M., Deorowicz, S., 2017. KMC 3: counting and manipulating k-mer statistics. Bioinformatics 33, 2759–2761.

Limasset, A., Rizk, G., Chikhi, R., Peterlongo, P., 2017. Fast and scalable minimal perfect hashing for massive key sets, in: Proc of SEA, pp. 25:1–25:16.

Marchet, C., Boucher, C., Puglisi, S., Medvedev, P., Salson, M., Chikhi, R., . Data structures based on k-mers for querying large collections of sequencing data sets. Genome Res 31, 1–12.

McVean, G., et al., 2012. An integrated map of genetic variation from 1,092 human genomes. Nature 491, 56–65.

Medvedev, P., Pham, S., Chaisson, M., Tesler, G., Pevzner, P., 2011. Paired de Bruijn graphs: A novel approach for incorporating mate pair information into genome assemblers. J Comput Biol 18, 1625– 1634.

Muggli, M., Alipanahi, B., Boucher, C., 2019. Building large updatable colored de Bruijn graphs via merging. Bioinformatics 35, i51–i60.

Muggli, M., Bowe, A., Noyes, N., Morley, P., Belk, K., Raymond, R., Gagie, T., Puglisi, S., Boucher, C., 2017. Succinct colored de Bruijn graphs. Bioinformatics 33, 3181–3187.

Munro, I., Nekrich, Y., Vitter, J.S., 2015. Dynamic data structures for document collections and graphs, in: Proc of PODS, pp. 277–289.

Noyes, N., et al., 2016. Resistome diversity in cattle and the environment decreases during beef production. eLife 5, e13195.

Overmars, M.H., van Leeuwen, J., 1981. Worst-case optimal insertion and deletion methods for decomposable searching problems. Inf Process Lett 12, 168–173.

Pandey, P., Almodaresi, F., Bender, M., Ferdman, M., Johnson, R., Patro, R., 2018. Mantis: A fast, small, and exact large-scale sequence-search index. Cell 7, 201–207.

Peng, Y., Leung, H.C., Yiu, S.M., Chin, F.Y., 2010. IDBA–a practical iterative de Bruijn graph de novo assembler, in: Proc of RECOMB, pp. 426–440.

Peng, Y., et al., 2012. IDBA-UD: A de novo assembler for single-cell and metagenomic sequencing data with highly uneven depth. Bioinformatics 28.

Pevzner, P., Tang, H., Waterman, M., 2001. An Eulerian path approach to DNA fragment assembly. Proc Natl Acad Sci 98, 9748– 9753.

Prezza, N., 2017. A framework of dynamic data structures for string processing, in: Proc of SEA, p. 11:1–11:15.

Ronen, R., Boucher, C., Chitsaz, H., Pevzner, P., 2012. SEQuel: improving the accuracy of genome assemblies. Bioinformatics 28, i188–i196.

Sirén, J., Garrison, E., Novak, A.M., Paten, B., Durbin, R., 2020. Haplotype-aware graph indexes. Bioinformatics 36, 400–407.

Turnbull, C., et al., 2018. The 100,000 genomes project: bringing whole genome sequencing to the nhs. Br Med J 361.

Turner, I., Garimella, K., Iqbal, Z., McVean, G., 2018. Integrating long-range connectivity information into de Bruijn graphs. Bioinformatics 34, 2556––2565.

Zerbino, D., Birney, E., 2008. Velvet: Algorithms for de novo short read assembly using de Bruijn graphs. Genome Res 18, 821–829.

